# Skilled strategies of release parameters for accurate free-throw shooting in the presence of motor noise

**DOI:** 10.1101/793513

**Authors:** Nobuyasu Nakano, Yuki Inaba, Senshi Fukashiro, Shinsuke Yoshioka

## Abstract

How humans execute accurate movement in the presence of motor noise is a key problem in the field of biomechanics and motor control that limits the performance improvement in daily or sporting activities. The aim of this study was to clarify the strategy of basketball players during free-throw shooting. Two possible hypotheses were examined: the players minimize the release speed to decrease signal-dependent noise or the players maximize the shot success probability by accounting for their variability. Eight collegiate players and one professional player participated in this study by attempting shots from the free-throw line using a motion capture system. The solution manifold consisting of ball parameters at release was calculated and the optimal strategy was simulated by considering ball parameter variability; this result was compared with the actual data. Our results showed that participants selected the solution of near-minimum release speed. The deviation of the measured release angle from the minimum-speed angle was close to zero (2.8 ± 3.1°). However, an increase in speed-dependent noise did not have a significant influence on the ball landing position through simulation. Additionally, the effect of release angle error on the ball landing position was minimum when using the minimum speed strategy. Therefore, the players minimize the release speed to minimize the effect of the release error on performance, instead of minimizing the speed-dependent noise itself. In other words, the strategy is “near-minimum-speed strategy” as well as “minimum-error-propagation strategy”. These findings will be important for understanding how sports experts deal with intrinsic noise to improve performance.

## 1. Introduction

Motor invariance of goal-directed movements is desired to improve our performance in daily activities and sports. However, the noise that randomly disturbs signals in the nervous system results from various processes. For example, motor noise is attributed to the physiological organization of the pool of motor neurons and their muscle fibers (Faisal et al., 2008). Owing to this, humans cannot completely reproduce the same movements even in simple tasks such as isometric force production (Jones et al., 2002) or arm reaching (Gordon et al., 1994; van Beers, 2003). Furthermore, the magnitude of this motor noise produced by a skeletal muscle is proportional to that of the average force produced by the muscle; thus, it is termed signal-dependent noise (Jones et al., 2002; Hamilton et al., 2004). It has been suggested that humans minimize the motor variability (*e.g.* endpoint of arm reaching) in the presence of such intrinsic noise (Harris & Wolpert, 1998; Todorov & Jordan, 2002; van Beers, 2009).

Although traditional studies on movement variability evaluated the performance using an error or standard deviation, recent studies have attempted to determine more than a pure outcome such as an error or standard deviation. Some studies have focused on the distributional aspects of execution variables with respect to goal variables (*e.g.* joint angles with respect to an end-point position in an arm-reaching). Pioneering work of uncontrolled manifold (UCM-) analysis (Scholz & Schöner, 1999; Latash et al., 2002) and its extensive work of goal equivalent manifold (GEM-) analysis (Cusumano & Cesari, 2006; Cusumano & Dingwell, 2013; John et al., 2016) have quantitatively decomposed the fluctuations in the execution space into task-relevant and task-irrelevant components. Sternad and collegues developed the solution manifold (*i.e.* a set of abundant solutions to achieve a motor task) and tolerane, noise, covariation (TNC-) analysis (Müller & Sternad, 2004; Cohen & Sternad, 2009) to differentiate between three different contributions to the motor task performance. Other studies on movement variability focused on the strategy to maximize the expected performance by modelling some form of the individual variability. These studies examined decision-making strategy under risks and rewards during rapid pointing (Trommershäuser et al., 2003, 2005) or other tasks (Nagengast et al., 2011; Tibshirani et al., 2017; Ota et al., 2016). These studies provide perspectives for a better understanding of how to improve performance in the presence of intrinsic noise. The present study complements and extends these works using a more practical motor task performed by well-trained participants; most of these studies considered the participant’s movement after training for a few hours or days as skilled movement.

Expert sports movement is useful for analyzing strategies to improve performance. Davids and collegues proposed that movement models from sports reveal fundamental insights into motor coordination and control (Davids et al., 2005). This is probably because the player expertise is acquired through long-term deliberate practice where training is focused on improving movement-specific performance (Ericsson, 2008; Seifert et al., 2013). For example, basketball players practiced at throwing multiple shots from a free-throw line (Keetch et al., 2005, 2008; Breslin et al., 2010, 2012). Although several studies on the movement variability in basketball shooting have examined the joint coordination from the dynamical system perspective (Button et al., 2003; Robins et al., 2006; Mullineaux & Uhl, 2010), these studies could not explain the control strategy at the execution level that contributed to the performance at the goal level. Therefore, in this study, we focus on the release strategy of basketball players during free-throw shooting to evaluate how sports experts execute an accurate aimed throwing in the presence of intrinsic noise.

Two hypotheses are proposed as the strategy for accurate aimed throwing. One involves minimizing the release speed that is required for hitting a target when throwing a projectile because minimizing the force imparted on the ball or energy expenditure to generate the release speed were believed to decrease motor variance (Brancazio, 1981; Miller & Bartlett, 1996; Robins et al., 2006); this can also be understood as minimizing the release speed decreases the signal dependent noise. However, this results in a correspondingly lower margin for error in basketball shooting (*i.e.*, a shallow trajectory reduces the entry area), thus the signal-dependent noise and the allowable margin between a ball and hoop are in a trade-off relationship (Bartlett et al., 2007). The other hypothesis involves maximizing the expected performance. In a previous throwing task, performers were found to choose the solutions that were robust against execution noise; *i.e.*, less sensitive to error or perturbations in the movement, instead of minimizing release speed (Sternad et al., 2011). Moreover, a simulation study that quantified the effect of execution error on the performance result in overarm or underarm aimed throwing tasks provided theoretical evidence for the existence of an optimum speed (Venkadesan & Mahadevan, 2017), which was observed during actual throwing movements (van den Tillaar & Ettema, 2003; Freeston et al., 2007). Therefore, through a thorough analysis of these two hypotheses, this study aims to reveal the strategy by which skilled players perform accurate free-throw shooting in order to improve throwing performance.

## 2. Method

### Participants

Eight male collegiate basketball players participated in experiment 1. The free-throw shot probability during the experiment for the participants in experiment 1 was 65 ± 15% (*c.f.* expert players in the study of Okazaki & Rodacki (2012): 62 ± 12% and under 18’s national team player in the study of Button et al. (2003): 67%). The mean (± standard deviation) age, height, and body mass of the participants were 19.4 ± 1.1 years, 176 ± 8 cm, and 70.5±4.3 kg, respectively. In addition, one male professional player participated in experiment 2 to measure top-level performer; his shot probability during the experiment was 99%. The participants provided written informed consent prior to the commencement of the study and the experimental procedure used in this study was approved by the Ethics Committee of the Graduate School of Arts and Sciences of the University of Tokyo for experiment 1 and by the Human Subjects Committee of the Japan Institute of Sports Sciences for experiment 2.

### Data collection

The experimental setup is shown in Figure 1a. Participants made 50 shots in experiment 1 and 100 shots in experiment 2 from the free-throw line after warm-up. The participants were instructed to shoot without using the backboard to score a basket. A total of 6-12 reflective markers were attached onto the ball. The coordinates of the reflective markers attached onto the participants’ bodies and the ball were recorded using a 16-camera motion capture system (Motion Analysis Corp, Santa Rosa, CA, USA) at a sampling rate of 200 Hz in experiment 1 and using a 20-camera motion capture system (VICON MX series, Vicon Motion Systems Ltd., Oxford, UK) at a sampling rate of 500 Hz in experiment 2.

**Figure 1:**
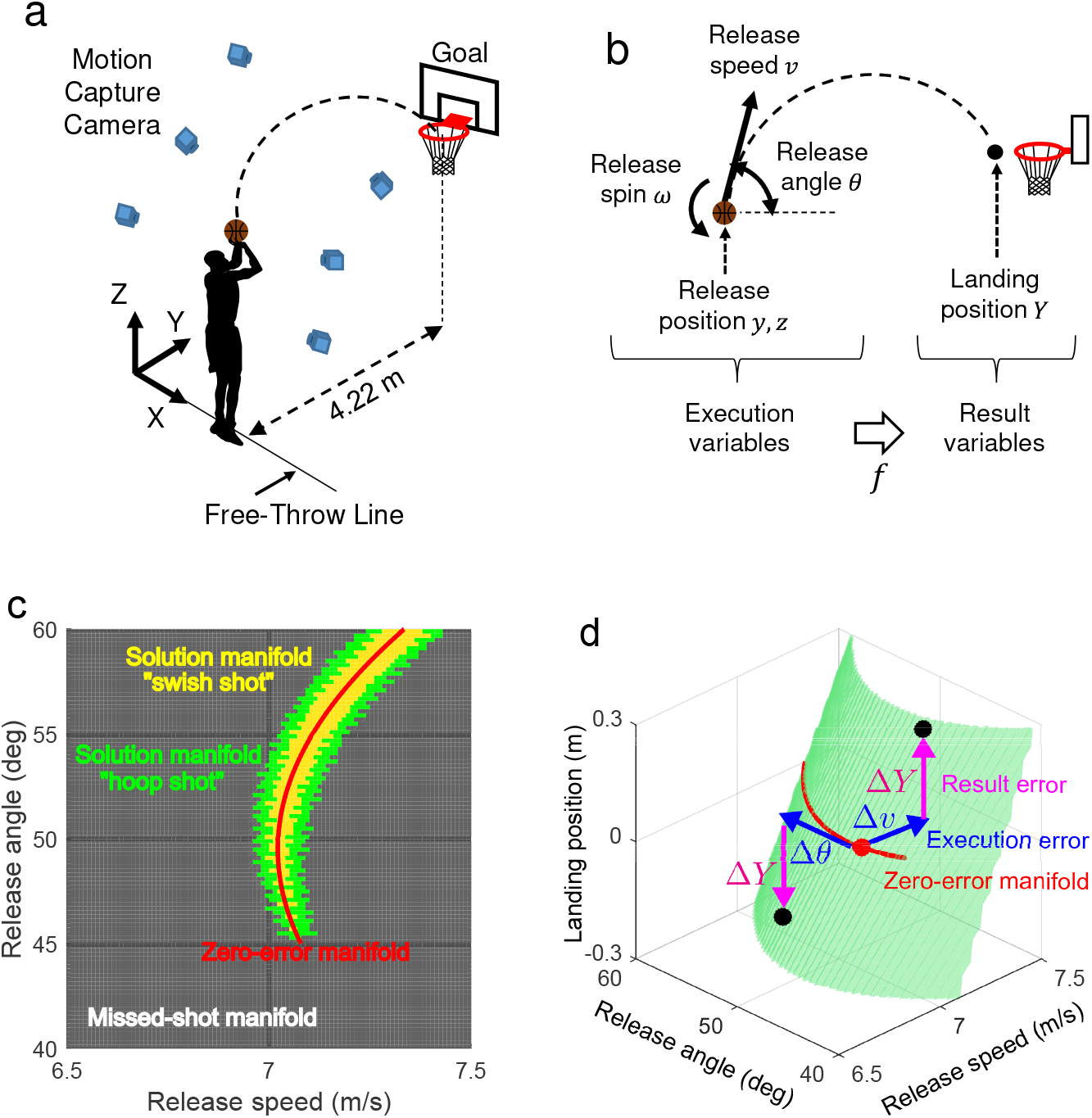
The experimental setup and the schematic description of the analysis. (a) The experimental setup of the free-throw shooting task. (b) The execution variables are defined as the ball release speed, angle, position, and spin in a two-dimensional plane. The result variables are defined as the ball landing position at the height of the hoop (= 3.05 m). A set of result variables is determined from a set of execution variables by calculating the trajectory. (c) A solution manifold in an execution space which is spanned by the release speed and angle. Yellow central area represents a set of variables which result in a swish shot (*i.e.* successful shot without contacting with hoop) based on (Brancazio, 1981), and green surrounding area represents a set of variables which result in a hoop shot (*i.e.* possibly successful shot by contacting with hoop) as defined by the landing at ±5 cm relative to the swish shot as in the study of Inaba et al. (2017). (d) The result error Δ*Y* caused by the execution error Δ*θ*, Δ*v* from a zero-error startegy. The magnitude of the gradient indicates how the execution error propagates to the performance result.

### Overview of data analysis

We focused on the relationship between the ball parameters at release (execution variables) and the ball landing position at the height of the hoop (result variables) (Figure 1b). In this study, we focused on the movements in the sagittal plane because the control of joints or ball in the plane is more important to performance for experienced players (Motoyasu et al., 2011; Hopla, 2012), as in the previous studies on basketball shooting (Miller & Bartlett, 1996; Button et al., 2003; Robins et al., 2006; Mullineaux & Uhl, 2010). First, we obtained the ball parameters at release from the measured coordinates of the reflective markers. Second, we identified the solution manifolds (*i.e.* a set of abundant solutions to achieve a motor task, (Müller & Sternad, 2004; Cohen & Sternad, 2009)) of the execution variables using simulations of the ball trajectory (Figure 1c) then compared the measured execution variables with the solution manifolds. Finally, to interpret the choice of execution strategy, we performed two different simulations: one estimated the shot probability according to the variability of each participant and the other quantified the effect of execution noise on the result (Figure 1d). Details of the data analysis are presented below.

### Release parameters

The ball release parameters (*i.e.* release speed, angle, spin, and position) were defined from the center of ball at the time of release. The details of the calculation for experiment 1 and 2 are described in the studies of Nakano et al. (2018) and Inaba et al. (2017), respectively.

### Simulation of a ball trajectory

To ensure the best match between the predicted ball trajectory and the measured trajectory, we used the equation of motion during flight including air force (Goff, 2013) as defined by Eq. (1). Here, ***f***_*D*_ and ***f***_*L*_ are the drag and lift force, which are defined in Eq. (2) (Goff, 2013), where *ρ* is air density, and *A*, ***v***, and ***ω*** are the cross-sectional area, velocity vector, and angular velocity vector of the ball. The trajectory was calculated by integrating Eq. (1) using the fourth-order Runge-Kutta method, in which time step size was set to 0.005 s. The two air coefficients (*C_D_*, *C_L_*) were estimated to fit the measured ball trajectories using the least squared method by assuming the coefficients to be constant, as in the study of Alaways & Hubbard (2001).

### Relative effects of release parameters on performance (preliminary analysis for the following analysis)

The landing position *Y* at the height of the hoop (goal) for a set of execution variables ***x*** was calculated as defined by Eq (3). The mean landing position 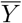 for the mean execution variables 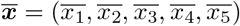 was calculated using by Eq (4). When *x*_*i*_ becomes *x*_*i*_ + Δ*x*_*i*_ and the other *x*_*j*_ (*j* ≠ *i*) are constant, that is, ***x*** increased by Δ***x***_*i*_ = [0, …, Δ*x_i_*, …, 0], the deviation of *Y* from the mean landing position 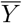 was calculated using Eq (5) and Eq (6). Here, we used the standard deviation of the measured release parameters as Δ*x_i_*. Thus, we defined the relative effects of the release parameters on the performance as Δ*Y* (Figure 2a). *y*, *z* and *ω* had a lower effect on the landing position compared with *θ* and *v*. In addition, the effect of *θ* increased considerably as the release angle increased (see Result, Figure 4). Therefore, in the following analysis, to simplify the problem and visualize the variables in the solution manifold, we considered the release speed *v* and release angle *θ* as variables, while the other release parameters were assumed to be constant as the mean value of each participant. This notion is also supported by the underhand throwing study (Dupuy et al., 2000).

**Figure 2:**
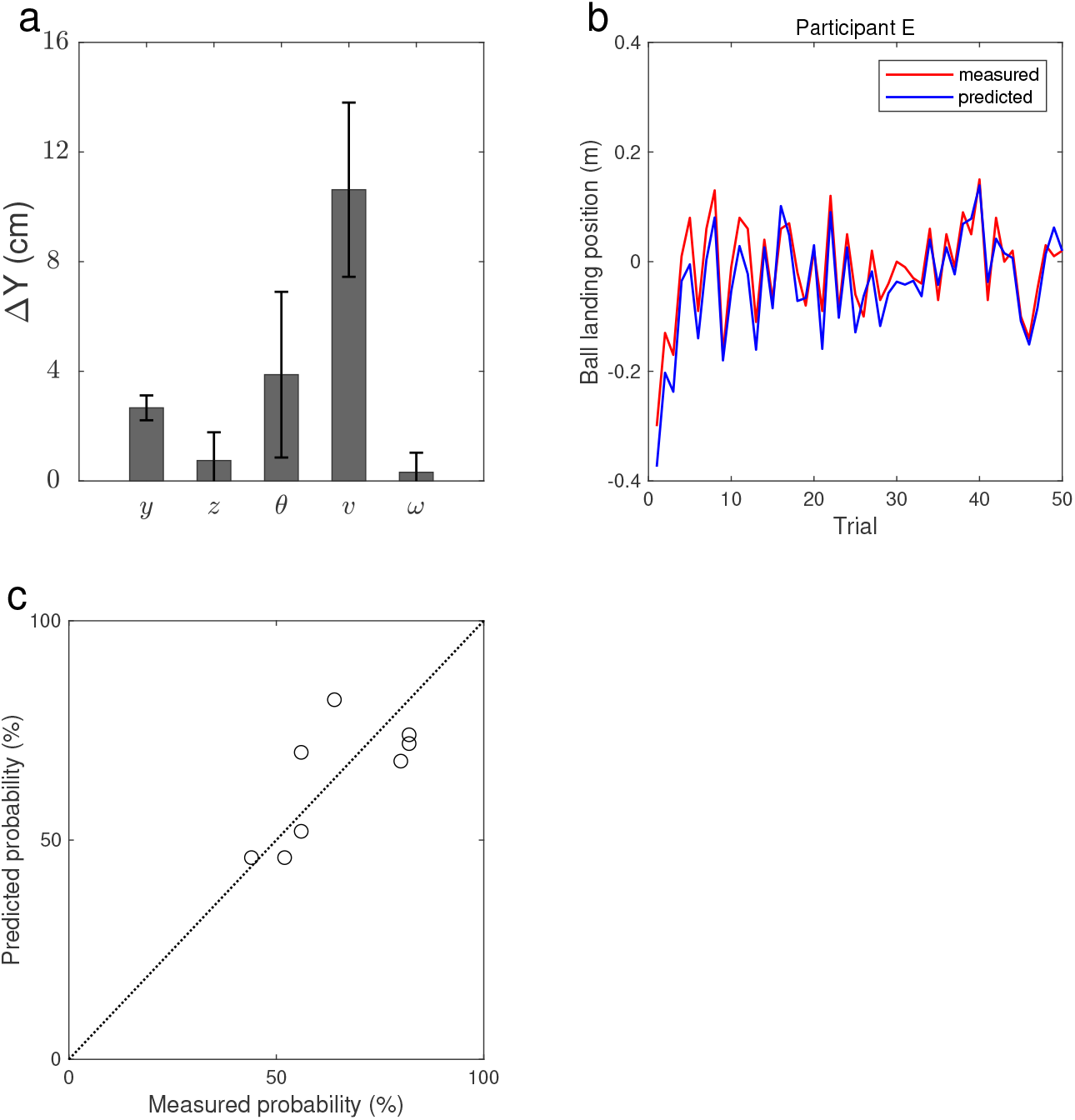
The results of the preliminary analysis to validate the simulation analysis. (a)Effects of release parameters on the landing position. The following results were calculated by assuming two release parameters (*i.e.* release speed and angle) as variables and the other parameters as constant. (b)Measured and predicted ball landing position of 50 trials for representative participant (participant E). Zero indicates the position of the hoop center. (c)Measured and predicted shot probability of each participant.

### Execution variables and solution manifold

For all combinations of release parameters, the results of the trajectories can be calculated using Eq. (1). The combinations of release parameters required to produce successful shots can be drawn as a solution manifold in an execution space (Müller & Sternad, 2004; Cohen & Sternad, 2009; Sternad et al., 2011). Two ranges of release speed and release angle were set to illustrate the execution variables of all participants. The local ranges of release speed and release angle ranged from 6.5-7.5 m/s and from 40-60°, and the global ranges ranged from 6-10 m/s and from 40-80°, respectively. Thus, the ranges were divided into 100 equal widths (*i.e.* 101 × 101 space). For each (*v*_*i*_, *θ*_*j*_) (*i* = 1, 2, …, 101, *j* = 1, 2, … 101), the success or miss of the simulated shot was determined based on the study of Brancazio (1981), in which the margin of error between the ball and the hoop was formulated for a given entry angle. A solution manifold in an execution space spanned by the release speed and angle is shown in Figure 1c. The central yellow area represents the set of variables that result in a swish shot (*i.e.* a successful shot without contacting with the hoop) based on Brancazio (1981). The green surrounding area represents the set of variables that result in a hoop shot (*i.e.* a possibly successful shot that contacts with the hoop), defined by the shot landing ±5 cm relative to the swish shot, as in the study of Inaba et al. (2017). The curved red line represents the set of variables that result in a zero-error (*i.e.* landing in the center of the hoop).

### Prediction of shot probability

To identify the optimal strategy, the probability of success was estimated using a statistical distribution. For each (*v*_*i*_, *θ*_*j*_) (*i* = 1, 2, …, 101, *j* = 1, 2, … 101), *N* random trials were generated from the bivariate normal distribution, where their probability density functions are given in Eq. (7). Here, 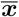 is the mean vector of ***x*** = (*v, θ*)^*T*^ and **Σ**_*i,j*_ is the covariation matrix of ***x***= (*v, θ*)^T^. We modeled the variance of the release parameters as speed-dependent noise (*i.e.* the variance is proportional to the speed) in Eq. (8). Here, **Σ**_0_ is the covariation matrix of measured data. The probability, *p*(*v*_*i*_, *θ*_*j*_), was calculated by computing the number of successful shots from *N* random trials, which was set to be 1000.

### Propagation of execution error to result error

To identify how the execution error propagated to the performance result for each strategy (*e.g.* how the error of 1° or 0.1 m/s in a strategy affects the landing position (Figure 1d)), we defined the index of error propagation. The predicted landing position of a ball at the height of the hoop can be calculated using Eq. (9). The small error in the landing position, *δy*, related to the error in the release parameter was estimated using linear approximation as defined in Eq. (10), where, 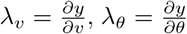, and the Jacobian, ***J***, is computed in the neighborhood of the zero-error manifold (*i.e.* the center of the solution manifold shown by the curved red line in Figure 1d) to examine the error propagation at various release angles. Because larger values of 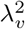 and 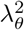 amplify the error of the execution variables, 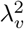 and 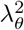 are considered to indicate the inaccuracy of the task (Venkadesan & Mahadevan, 2017). To match the units of both *v* and *θ* terms in this study, we adopted the index of error propagation that indicates the effect of execution error on the landing position, as defined by Eq. (11), where, we set Δ*v* and Δ*θ* to the standard deviations of *v* and *θ* for each participant.

## 3. Results

Figure 2b illustrates the measured and predicted ball landing position of 50 trials for representative participant (participant E). These results show that the predicted ball landing positions correspond to the measured ball landing positions. The mean absolute error of the ball landing position for all participants was 0.040 ± 0.019 m (4.0 ± 1.9 cm). Figure 2c illustrates the measured and predicted shot probability for all participants. Here, the predicted shot success included both the “swish shot” (yellow manifold in Figure 1c) and the “hoop shot” (green manifold in Figure 1c). The results show that the predicted shot probability correspond to the measured (*c.f.* Pearson’s correlation coefficient: *r* = 0.70, *p* = 0.05). Therefore, the ball trajectory simulation assumed two release parameters (*i.e.* release speed and angle) as variables and the other parameters as constant.

Figure 3 illustrates the execution variables of all trials and solution manifolds of the representative participants in the form of global and local ranges of variables. Note that the solution manifolds were drawn for each participant, and they were not the same. Regarding the distribution of execution variables in the global ranges (Figure 3a, c, and e), all participants preferred the strategy of near-minimum release speed to that of higher release speed. Furthermore, the average deviation of the mean angle, 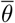, from the minimum-speed angle, 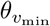, was close to zero (2.8 ± 3.1°) for all participants, and the maximum deviation was 7.2°. None of the participants used the strategies with angles over 60° (*i.e.* higher trajectories).

**Figure 3:**
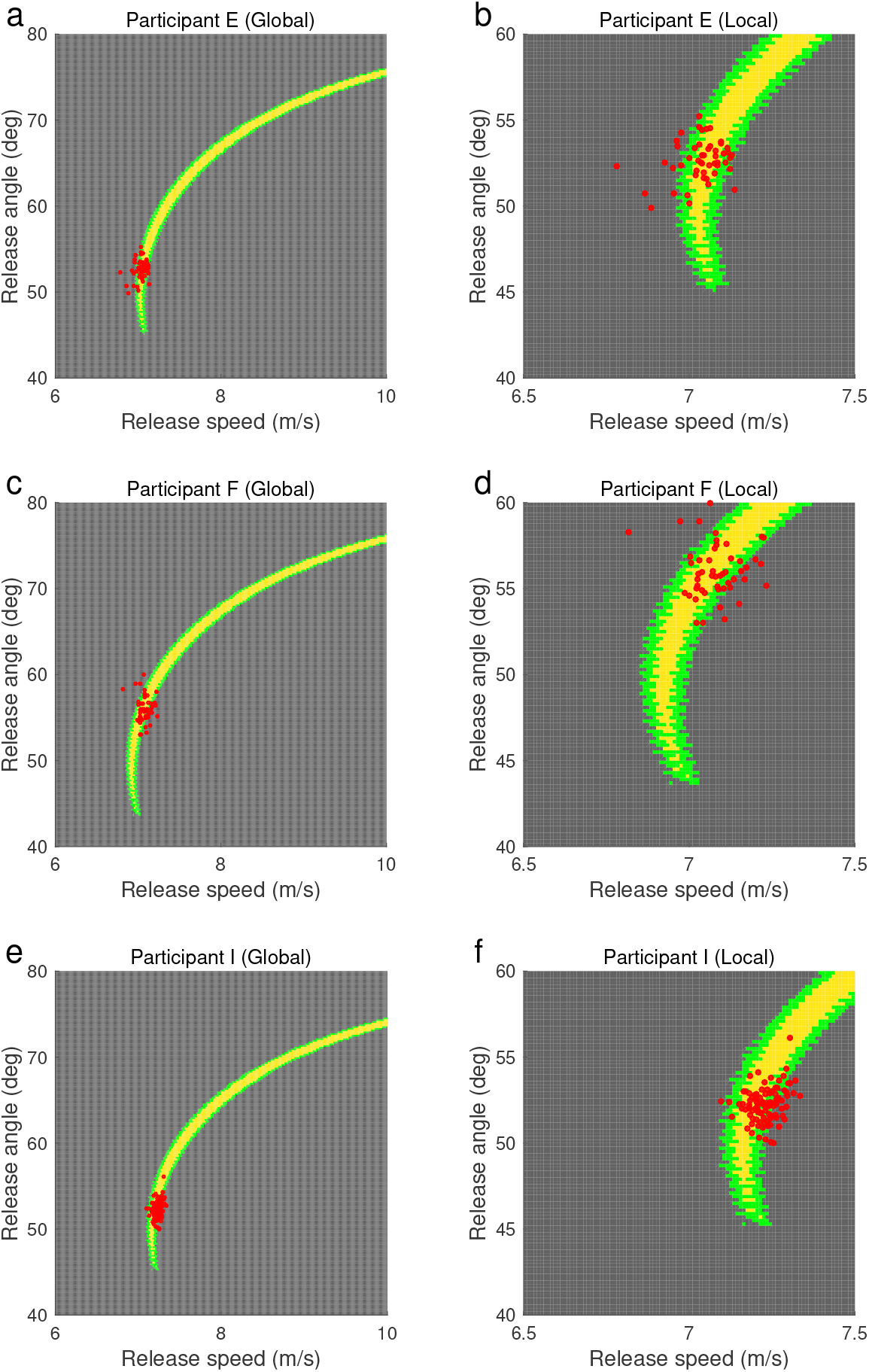
Execution variables and solution manifolds of each participant. Left and right panels show the results in the wide and narrow range of release speed and release angle, respectively. Yellow central area represents a set of variables which result in a swish shot (i.e. successful shot without contacting with hoop) based on (Brancazio, 1981), and green surrounding area represents a set of variables which result in a hoop shot (i.e. possibly successful shot by contacting with hoop) as defined by the landing at ±5 cm relative to the swish shot as in the study of Inaba et al. (2017). Red filled circles represent actual execution variables of each participant.

In addition, we calculated the index of error propagation (*λ*_*v*_ · Δ*v*)^2^ + (*λ*_*θ*_ · Δ*θ*)^2^ to explain the advantages of using the strategy of near-minimum release speed. Figure 4 shows that the error propagation of the release angle was smallest (*i.e.* least sensitive to angle error) at the minimum-speed angle (49.6° for this participant); this then rapidly increased (*i.e.* became more sensitive to angle error) as the deviation of the release angle from the minimum-speed angle increased. Figure 4 also shows that the error propagation of the release speed decreased monotonically and gradually (*i.e.* became less sensitive to speed error) as the release angle increased. Thus, the index of error propagation (*λ*_*v*_ · Δ*v*)^2^ + (*λ*_*θ*_ · Δ*θ*)^2^ was the smallest at the near-minimum-speed angle (51.2° for this participant). Therefore, using the strategy of near-minimum release speed made the performance result less sensitive to execution error.

**Figure 4:**
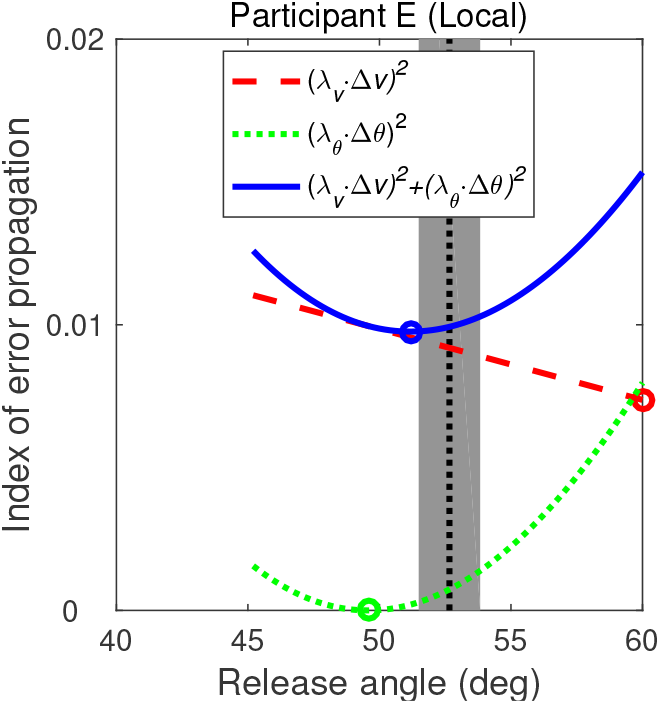
The index of error propagation that indicates the effect of the execution error on the landing position. Red dashed line and green dotted line indicate the individual effect of error in *v* and *θ*. Blue solid line indicates the index of error propagation. Circle indicates the minimum value for each line. Black dotted line and gray filled area indicate the mean and standard deviation of the release angle for each participant.

Figure 5 illustrates the simulated probability of success for each (*v*_*i*_, *θ*_*j*_) in the form of the global and local ranges of variables. The optimal strategy to achieve the highest probability was found to be the solution at the angle (60.8° for the participant) that was larger than the minimum-speed angle. The optimal release angles required to achieve the highest probability tended to be larger than the actual release angles; this was especially true for the participants who exhibited higher shot probability.

**Figure 5:**
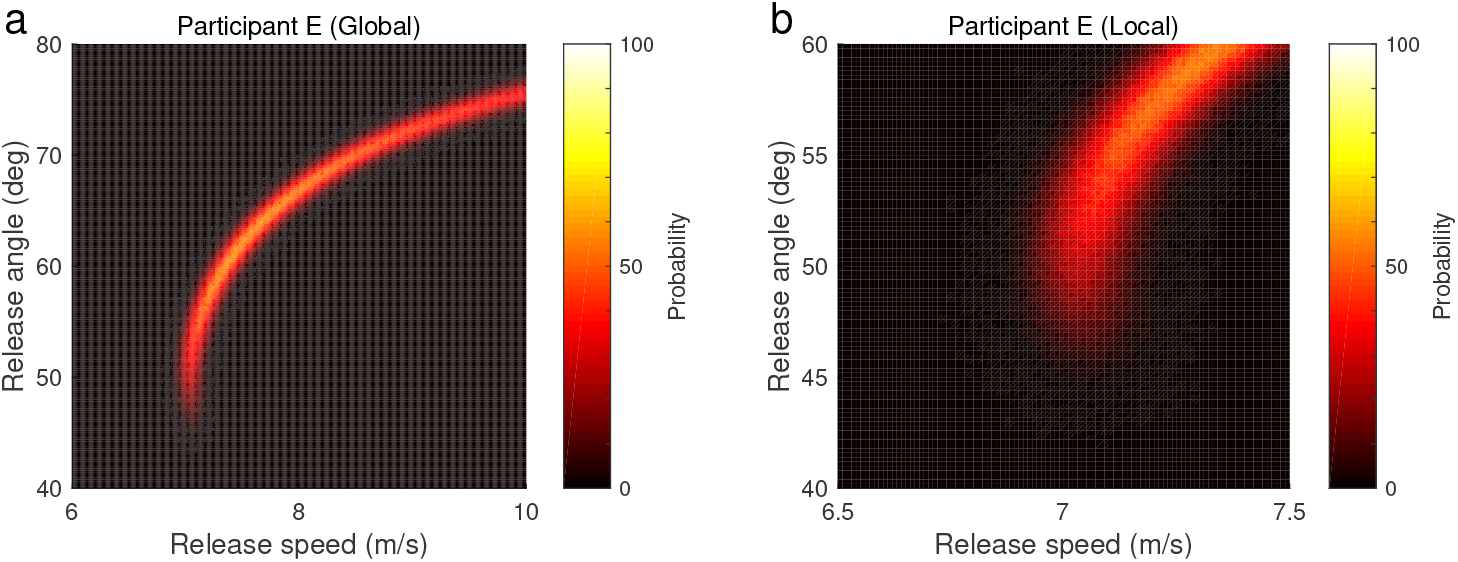
Simulated shot probability of success using speed-dependent covariant noise model. Left and right panels show the results in the wide and narrow range of release speed and release angle, respectively.

Regarding the distribution of execution variables in the local ranges (Figure 3b, d, and f), the chosen strategies were not the absolute minimum-speed. The deviation of the mean angle, 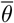, from minimum-speed angle, 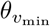, for each participant is shown in Table 1. The deviation between the measured strategy and the minimum-speed strategy for each participant ranged from −2.1° to 7.2°. Most participants selected an angle that was slightly larger than the minimum-speed angle.

**Table 1:** Measured shot probbility of success in the experiment and the deviation of the mean angle 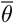 from minimum-speed angle 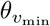 for each participant

| Participant | Probability (%) | $\bar{\theta} - \theta_{v_{\min}}$ (deg) |
| --- | --- | --- |
| A | 82 | -2.1 |
| B | 52 | 0.5 |
| C | 44 | -0.9 |
| D | 56 | 5.3 |
| E | 82 | 3.3 |
| F | 56 | 7.2 |
| G | 64 | 5.4 |
| H | 82 | 3.5 |
| I | 99 | 3.1 |
| Mean±SD | | 2.8±3.1 |

## 4. Discussion

The aim of this study was to elucidate the strategy used by experts to improve their performance in free-throw shooting. We therefore tested two hypothesized strategies, whereby players minimized the release speed to decreases the signal dependent noise or selected appropriate release parameters to maximize the shot probability.

### Hypothesis: players minimize release speed

The minimum-speed strategy was proposed in previous studies on basketball shooting (Brancazio, 1981; Miller & Bartlett, 1996; Robins et al., 2006). The factors of the strategy have been recognized to minimize energy expenditure; however, they have not been quantitatively described in previous studies. In this study, without using the concept of energy expenditure, which cannot be quantified, we examined the factors that determine the strategy through simulation by using a signal dependent noise model.

All participants used the strategy of near-minimum release speed according to the global ranges of execution variables (Figure 3); this is consistent with the studies of underarm precision throwing task imitating a *petanque*, in which players throw a projectile towards a target on the ground at the distance of 4-8 m (Dupuy et al., 2000). At first glance, this seems to be inconsistent with the previous studies where participants chose error-tolerant solutions (*i.e.* high probability) rather than the minimum-speed solution in a virtual throwing task imitating a *skittles* game (Sternad et al., 2011). This inconsistency may be due to the different shapes of the solution manifolds for each task. The solution manifolds of this free-throw task indicate that the error propagation of the release angle was minimized by using the angle of near-minimum speed (Figure 4); this made the performance insensitive to the error of the release angle when choosing the strategy of near-minimum speed. Conversely, the solution manifolds of virtual *skittles* tasks (experiment 2 in Sternad et al. (2011)) indicate that the error propagation of the release speed was much smaller than that of the release angle; this allowed participants to freely choose any release speed. For both studies, it is likely that players care less about adjusting the parameter that has a smaller error effect on the performance and focus instead on adjusting the other parameter.

The strategy of near-minimum speed may be advantageous when players choose a successful ball trajectory from various distances. In the game situation, players need to shoot from various distances; thus, it would be difficult for players to calculate the optimal release angle by considering their variability at each location. Our previous study, where basketball players performed jump shots from three different distances, showed that the release speed differed significantly between distances, but the release angle did not (Nakano et al., 2018). It is possible that players focus on adjusting the release speed by using the release angle of near-minimum speed. Note that when the defender is very close to a shooter in a game, the shooter does not always use the strategy because they use a larger release angle in the presence of the defender to avoid the block from the defender (Rojas et al., 2000).

Further, speed-dependent noise did not have a substantial influence on performance within local ranges for the same task (Figure 5). This is because the local range of release speed, which was selected by the participants, was very small in the free-throw task. Additionally, the error propagation of the release speed decreased as the speed and speed-dependent noise increased (*i.e.* higher trajectory) (Figure 4). These results suggest that minimizing the release speed is related to minimizing the error propagation of the release parameters, rather than minimizing the speed-dependent noise itself.

### Hypothesis: players maximize shot probability

To predict the optimal strategy, we performed simulation analysis by modeling the measured variability of each participant in the experiments using a bivariate normal distribution, similar to the decision making studies for several tasks such as rapid pointing (Trommershäuser et al., 2003, 2005). Most of the participants in this study selected the strategy that was different from that of highest probability (Figure 3 and Figure 5), although human pointing under reward and risk was consistent with the strategy that would maximize the expected gain (Trommershäuser et al., 2003, 2005). The non-optimality based on individual preference or experience in motor control has been argued in recent studies. For example, the individual behavior in a speeded aiming movement was sub-optimal with respect to maximizing the expected gain and was in acccordance with a risk-sensitive account of movement selection (Nagengast et al., 2010, 2011); this was retained throughout extensive practice (Ota et al., 2016). Moreover, the individual internal model of individuals’ motor error in speeded reaching was inaccurate after training (Zhang et al., 2013, 2015). Therefore, it is likely that participants in our free-throw experiment did not have the accurate noise model that was assumed in this study.

### Integration of two hypotheses

At a first glance the results seem to support hypothesis 1 (Figure 3 and Figure 4). However, only one of the two hypotheses should not be rejected or accepted because the two hypotheses are related. In human movement studies, cost functions proposed for biological movements can generally be divided into effort (some form of activation) and variability (performance accuracy) costs. The relative importance of the two costs are dissociated and estimated in the bimanual force production task of O’Sullivan et al. (2009); however, these two costs are usually related and cannot be dissociated. It is important to understand whether the effort cost itself contributes to the performance, or whether the variability cost also contributes at the same time. In this study, the release speed was minimized due to decreasing the effect of release parameter errors on their performance, instead of decreasing speed-dependent noise itself. In other words, the strategy used by the participants is “near-minimum-speed strategy” as well as “minimum-error-propagation strategy”.

### Limitations

The strategy of skilled basketball players was to use the angle of near-minimum speed, but not the absolute minimum speed in the local range (Figure 3b, d, f and Table 1). This difference may indicate a slight shift from the minimum speed strategy to the maximum probability strategy (Figure 5); however, this was unfortunately not clear from our results.

The noise model assumed in this study cannot be determined to be correct (*e.g.* the noise linearity in the release speed or distribution normality of release parameters). For a better understanding of the noise characteristic, an additional experiment is necessary, where more trials are performed for measuring the release speed and angle noise at various release speeds.

Note that this study was completely confined to the analysis of the ball release parameters. The kinematic or kinetic level of human movement is beyond the scope of this study. Our previous study, where basketball players performed jump shots from three different distances, suggested that the lower limb played a role in increasing or decreasing the energy output, and the upper limb played a role in compensating the variability of transferred energy to the ball (Nakano et al., 2018). Therefore, the causal mechanism behind accurate free-throw shooting or similar aimed movements at the level of kinematics or kinetics requires further clarification.

### Conclusion

Our results showed that most participants, especially the high-scoring participants, selected the solution of near-minimum release speed rather than the simulated optimal strategy based on their variability. Thus, we conclude that skilled players almost minimize the release speed to minimize the effect of release parameter errors on their performance, instead of minimizing speed-dependent noise itself. This suggests that players emphasize on adjusting the release speed by using the strategy that is robust to release error.

## Acknowledgements

This work was supported by JSPS KAKENHI Grant Number JP19J13288. The authors would like to thank Kazutoshi Kudo at the University of Tokyo for the helpful comments and members of the sports biomechanics laboratory at the University of Tokyo for the data collection in the experiment.

## Conflict of interest

There are no conflicts of interest to declare.

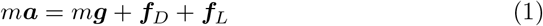

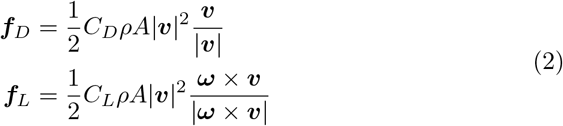

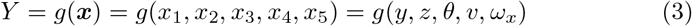

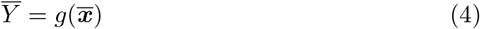

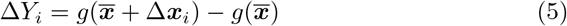

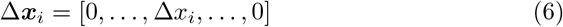

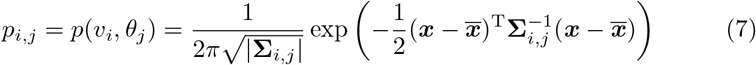

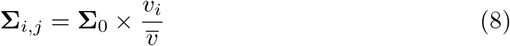

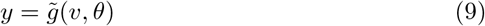

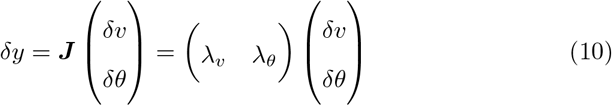

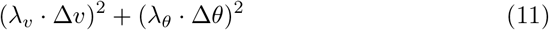

